# A slight decrease in serum albumin level is associated with rapid progression of kidney dysfunction even within the normal range: The Yuport Health Checkup Center Cohort Study

**DOI:** 10.1101/655563

**Authors:** Hoichi Amano, Kazunobu Yoshimura, Ryutaro Iijima, Kaito Waki, Keisei Matsumoto, Hitomi Ueda, Yasuko Ito, Ken Miyamoto, Kimihiko Akimoto, Takashi Yokoo, Kazuo Inoue, Hiroyuki Terawaki

## Abstract

**Objective:** A low-normal albumin level is associated with a high risk of cardiovascular disease and mortality in the general population. However, the relationship between serum albumin level and future decline of kidney function is unclear. We aimed to clarify the effect of serum albumin level on the decline of kidney function in the general population.

**Methods:** The data used were from 11,000 participants of a voluntary health checkup program between 1998 and 2006 conducted in Japan. The primary outcome for kidney function was a difference in estimated glomerular filtration rate (ΔeGFR) of ≥3 mL/min/1.73 m^2^/year. The association of the risk of decreased kidney function with albumin level was determined using a logistic regression analysis. We fit separate multivariable logistic regressions for serum albumin levels (g/dL) as a continuous variable and as categorical data, classified as ≤4.3 (n=2,530), 4.4– 4.6 (n=5,427), and ≥4.7 (n=3,043).

**Results:** Of 11,000 participants, 346 had a ΔeGFR/year of ≥3. As compared with the participants with albumin levels of ≥4.7 g/dL, the risk of decline in kidney function was higher not only in those with albumin levels of ≤4.3 g/dL (adjusted OR = 2,29, 95% CI: 1.65–3.18) but also in 4.4-4.6 g/dL (adjusted OR = 1.60, 95% CI: 1.20–2.14).

**Conclusion:** Decreased albumin level is an independent risk factor for rapid decline in kidney function even within the normal range.

## INTRODUCTION

The number of patients with chronic kidney disease (CKD) has been increasing in most parts of the world, and the disease is estimated to affect 200 million individuals worldwide [1]. Furthermore, the increase in the number of patients with CKD is expected to accelerate. CKD creates a large burden and is recognized as an important problem for both individuals and the society as a whole. First, CKD is a risk factor for not only end-stage kidney disease (ESKD) but also cardiovascular disease (CVD), which is the main cause of death worldwide [2-5]. Second, the worldwide medical expenses associated with hemodialysis due to ESKD is estimated to increase to a 1000 billion USD within the next 10 years [6]. For these reasons, establishment of an effective measure for CKD prevention is vital; in fact, this is one of the most important issues in public and national health.

While accumulating evidence shows that some metabolic and lifestyle risk factors of CKD, such as hypertension, dyslipidemia, and diabetes mellitus were addressed [7-13], the effective measure for CKD prevention has not been established yet. Therefore, risk factors other than the “conventional” risk factors of CKD should be considered.

Previous studies suggested that a lower albumin level, even that within the clinical normal range, is associated with high risks of CVD and mortality in the general population [14,15]. However, studies that investigate the relationship between the albumin level and decline of kidney function are completely lacking.

The aim of this study was to evaluate the effect of serum albumin level on the decline of kidney function in the general population by using a large retrospective cohort data set of the Japanese population.

## METHODS

### Study design and study population

This was a retrospective cohort study, and we used a dataset derived from the health screening program performed by the Yuport Medical Checkup Center in Tokyo. In this study, we set the 4-year baseline period to be between April 1998 and March 2002, and the 4-year follow-up period between April 2002 and March 2006. During the baseline period, 21,885 persons underwent checkups at least once during this period when, in total, 47,995 checkups were performed (Fig. 1). If the subjects underwent more than one checkup during the baseline period, the initial checkup data were used. During the follow-up period, 23,547 persons underwent checkups at least once for 49,390 checkups. If the subjects underwent more than one checkup during the follow-up period, all the data were used to identify incident diabetes. Follow-up data were merged with baseline data, yielding 11,129 persons who had been examined during both time periods. Of these patients, 129 with known diabetes at baseline were excluded, leaving 11,000 persons.

**Figure 1.**
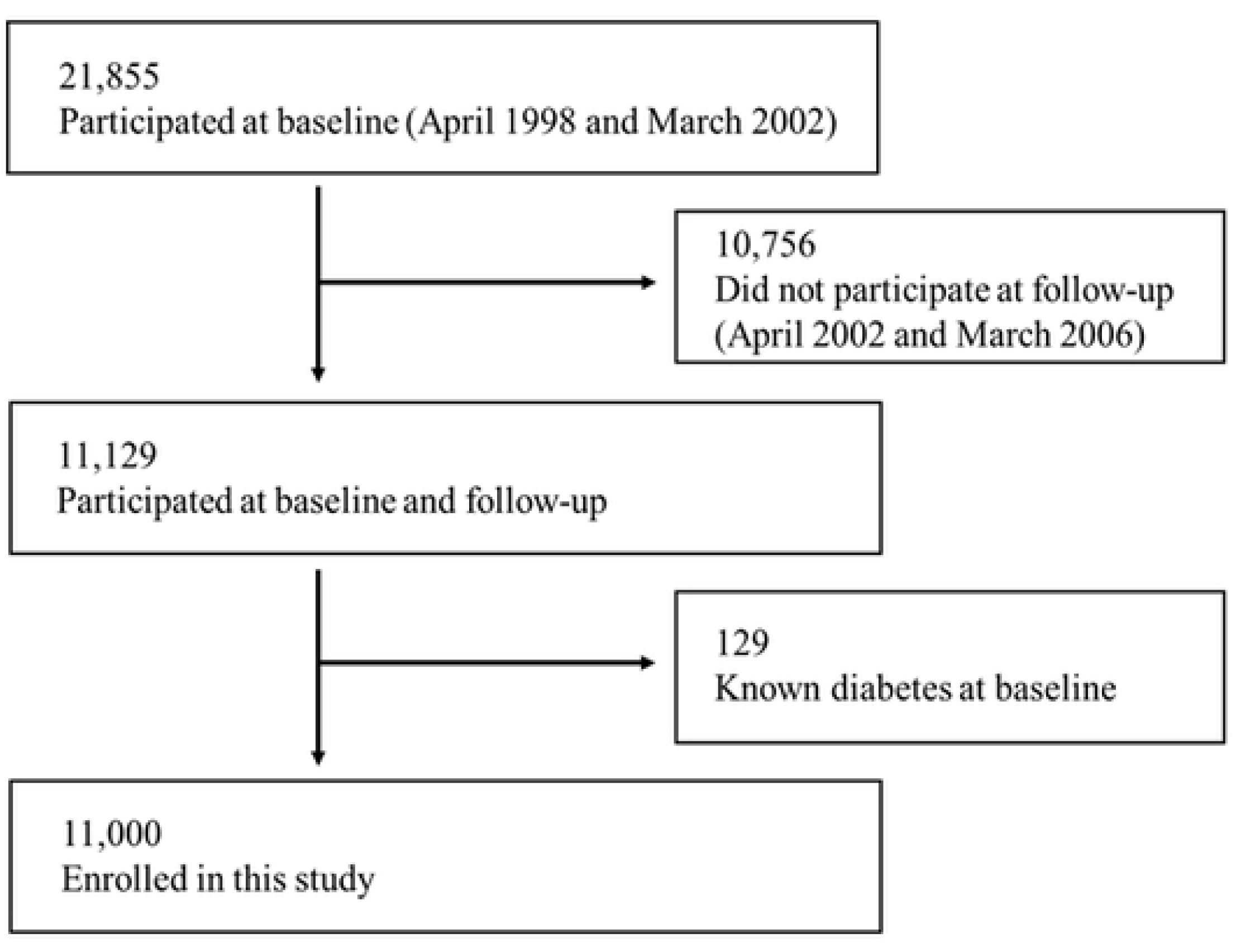
Flowchart of the study population.

In accordance with the Private Information Protection Law, information that might identify subjects was kept private by the center. Informed consent for anonymous participation in epidemiological research was obtained at every checkup.

### Measurements

Serum albumin level was determined using the bromocresol green method (reagents supplied by Denka Seiken, Tokyo, Japan) [16]. Other laboratory values were measured using the standard laboratory technique. All the checkup procedures were performed in the same manner, both during the baseline and follow-up periods, including blood measurements. Height and weight were measured to calculate body mass index (BMI), which was defined as weight divided by height squared (kg/m^2^). Blood pressure was measured by trained nurses using a sphygmomanometer.

### Kidney function

Kidney function was expressed as an estimated glomerular filtration rate (eGFR) using the CKD Epidemiology Collaboration (CKD-EPI) modified for Japanese [17]. GFR estimated by the coefficient-modified CKD-EPI equation was more closely related to CVD incidence than that estimated by the Japanese GFR equation [18]. The coefficient-modified CKD-EPI equation is as follows.

Estimated GFR (mL/min/1.73 m^2^) = 141 × min (Cr/κ, 1)^α^ × max (Cr/κ, 1)^-1.209^ × 0.993^Age^ × 1.018 (if female) × 0.813 (Japanese coefficient).

(κ: 0.7 in females and 0.9 in males, α: −0.329 in females and −0.411 in males.)

In general, kidney function decreases with age: A previous study showed that the rate of GFR decline was 0.36 ml/min/1.73 m^2^/year on average among 120,727 individuals aged ≥40 years [19]. Therefore, the clinical kidney disease outcome in this study was assessed as an abnormal annual decline of kidney function in each participant. An abnormal annual decline of kidney function was defined as a difference in eGFR (ΔeGFR) of ≥3 mL/min/1.73 m^2^/year. This cutoff value for an “abnormal decline,” represents a magnitude of change that is >3 times the rate previously described in studies of normal aging, and this change is known to be associated with clinically deleterious outcomes [20, 21]. To ensure the accuracy of evaluated trend, we also performed a sensitivity analysis in which a ΔeGFR of ≥5 mL/min/1.73 m^2^/year was regarded as an abnormal decline [22].

### Statistical analyses

Continuous data were expressed as mean ± standard deviation or median within the 25th and 75th percentiles, and categorical data were expressed as percentages. Baseline characteristics were compared between those who met the criteria for abnormal decline in kidney function and those who did not, and independent variables were assessed using the chi-square test in the case of categorical variables and the t-test or Mann-Whitney U test in the case of continuous variables. Analysis for trend was evaluated using Cochran-Armitage test. Regarding the relationship between two values, the Pearson correlation coefficient was employed.

Abnormal decline in kidney function and odds ratios (ORs) were estimated from the logistic regression model. Multiple analyses were used to calculate the OR for abnormal decline in kidney function after adjusting for age, sex, BMI, systolic blood pressure (SBP), eGFR at baseline, serum alanine aminotransferase level, serum uric acid level, high-density lipoprotein (HDL) level, HbA1c level, C-reactive protein level, history of cardiovascular disease (stroke or ischemic heart disease, or both). Inclusion of variables in the models was based on our existing knowledge regarding the risk factors of kidney function decline. We fit separate multivariable logistic regressions for both serum albumin level as a continuous variable and categorical data, which were classified as ≤4.3, 4.4– 4.6, and ≥4.7 g/dL.

Differences with a P value of <0.05 were considered statistically significant. All statistical analyses were performed using EZR Version 1.33 (Saitama Medical Center, Jichi Medical University, Saitama, Japan), which is a graphical user interface for R (The R Foundation for Statistical Computing, Vienna, Austria). More precisely, it is a modified version of R commander, which is designed to add statistical functions frequently used in biostatistics [23].

### Ethics issues

This study was conducted in accordance with the principles of the Declaration of Helsinki. The study was approved by the review board of Teikyo University (approval No. 15-205). The participants’ written informed consent for anonymous participation in epidemiological research was obtained at every evaluation.

## RESULTS

The baseline characteristics of the participants according to abnormal decline in kidney function status are shown in Table 1. Of the participants, 346 had an abnormal decline in kidney function. Among the patients with abnormal decline in kidney function, higher SBP values and lower levels of albumin and HDL-C were observed. The relationship between baseline serum albumin level and the change in eGFR is shown in Fig. 2.

**Table 1:** Participants’ characteristics at baseline according to decline in kidney function.

| | All | ΔeGFR/year < 3<br>mL/min/1.73 m<sup>2</sup><br>(n = 10,654) | ΔeGFR/year ≥ 3<br>mL/min/1.73 m<sup>2</sup><br>(n = 346) | P value |
| --- | --- | --- | --- | --- |
| Age (years) | 53.2 (11.6) | 53.2 (11.6) | 52.2 (13.4) | 0.104 |
| Sex (male), n (%) | 5248 (48) | 5101 (48) | 147 (43) | 0.049 |
| Body mass index (kg/m<sup>2</sup>) | 23.0 (3.0) | 23.0 (3.0) | 23.1 (3.2) | 0.387 |
| Systolic blood pressure (mmHg) | 124.1 (17.9) | 124.0 (17.9) | 126.5 (18.7) | 0.011 |
| Diastolic blood pressure (mmHg) | 75.0 (11.0) | 74.9 (11.0) | 77.5 (11.4) | <0.001 |
| **Laboratory data** |  |  |  |  |
| Hemoglobin (g/dL) | 13.9 (1.4) | 13.9 (1.4) | 13.9 (1.6) | 0.460 |
| Serum albumin (g/dL) | 4.52 (0.23) | 4.52 (0.23) | 4.46 (0.23) | <0.001 |
| Aspartate aminotransferase (U/L) | 22.9 (16.7) | 22.8 (16.7) | 22.2 (16.1) | 0.476 |
| Alanine aminotransferase (U/L) | 22.7 (9.3) | 22.69 (9.35) | 22.25 (8.70) | 0.393 |
| Lactate dehydrogenase (U/L) | 298.2 (55.3) | 298.1 (55.2) | 299.29 (57.5) | 0.701 |
| Blood urea nitrogen (mg/dl) | 14.9 (3.5) | 14.9 (3.5) | 14.9 (3.8) | 0.989 |
| Serum creatinine (mg/dL) | 0.7 (0.2) | 0.72 (0.16) | 0.73 (0.2) | 0.191 |
| Estimated glomerular filtration rate (mL/min/1.73 m<sup>2</sup>) | 82.9 (10.3) | 82.4 (10.2) | 83.8 (13.3) | 0.090 |
| Serum uric acid (mg/dL) | 5.4 (1.4) | 5.4 (1.4) | 5.4 (1.5) | 0.951 |
| Total cholesterol (mg/dL) | 203.3 (35.0) | 203.4 (34.9) | 201.0 (36.8) | 0.206 |
| High-density lipoprotein cholesterol (mg/dL) | 58.5 (15.2) | 58.6 (15.2) | 55.7 (14.2) | <0.001 |
| Triglyceride (mg/dL) | 97.0 [70.0, 140.0] | 97.0 [69.0, 140.0] | 97.5 [73.0, 143.0] | 0.308 |
| HbA1c (%; NGSP) | 5.1 (0.7) | 5.1 (0.7) | 5.1 (1.0) | 0.770 |
| C reactive protein (mg/dL) | 0.10 [0.01, 0.10] | 0.10 [0.01, 0.10] | 0.10 [0.01, 0.10] | 0.092 |
| **History of complications** |  |  |  |  |
| Stroke (+, %) | 20 (0.2) | 19 (0.2) | 1 (0.3) | 0.473 |
| Angina pectoris (+, %) | 36 (1.2) | 32 (0.3) | 4 (1.2) | 0.026 |
| Myocardial infarction (+, %) | 10 (0.1) | 10 (0.1) | 0 (0) | 0.999 |
| Observation period (years) | 5.4 (1.6) |  |  |  |
| Data are expressed as mean and standard deviation, or percentage and number or median with 25th and 75th percentiles. Abbreviations: NGSP, National Glycohemoglobin Standardization Program |  |  |  |  |

**Figure 2.**
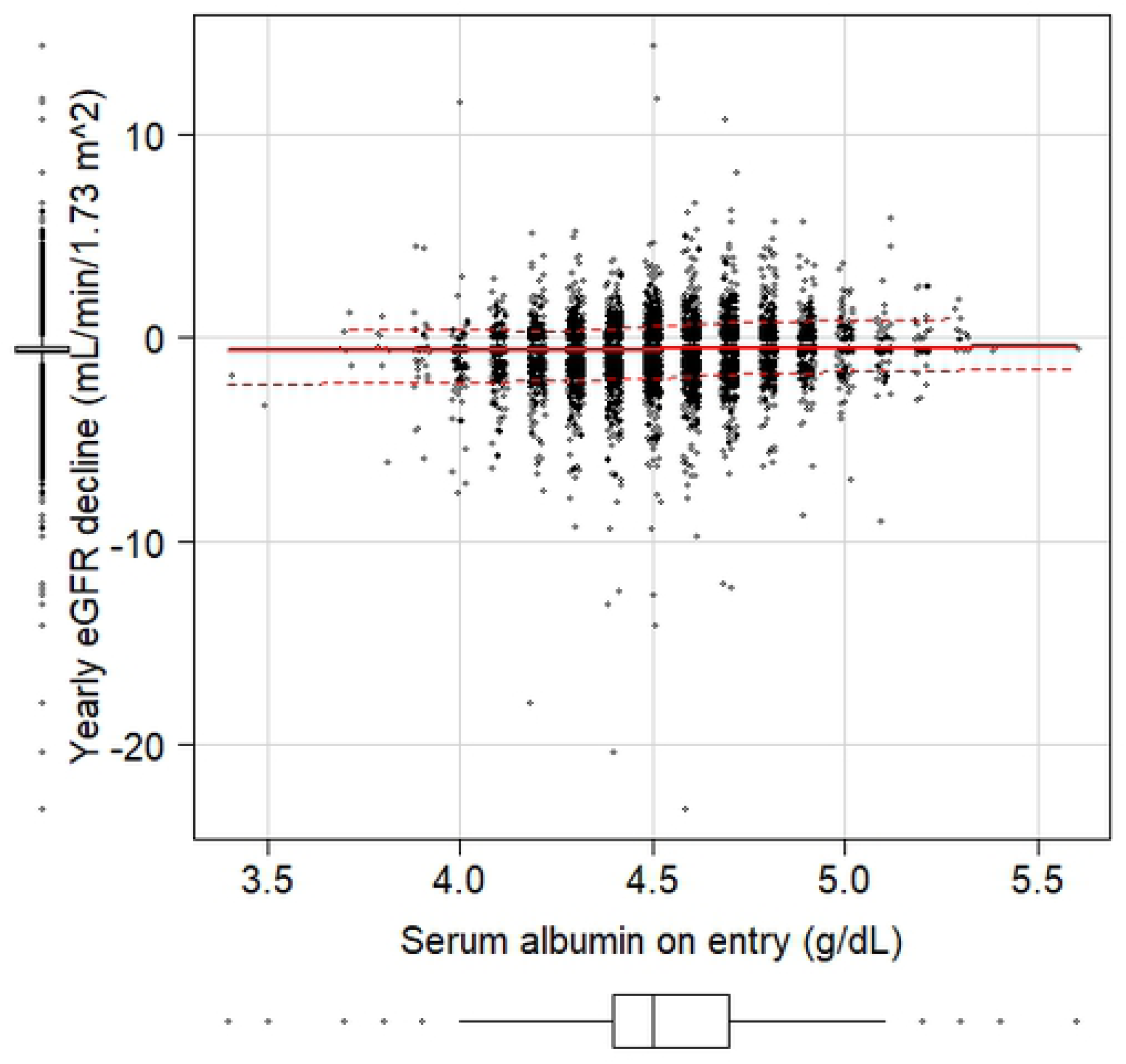
The relationship between baseline serum albumin level and the yearly decline in kidney function. A weak but significant relationship was observed (R = −0.0830, P = 2.92 × 10^-18^): Lower serum albumin level was related to greater decline in estimated glomerular filtration rate.

Next, we evaluated the unadjusted and adjusted ORs and 95% confidence intervals (CIs) for abnormal decline of kidney function according to albumin level. In the continuous variable evaluation, the albumin levels (per 0.1 g/dL) were negatively associated with the risk of abnormal decline in kidney function in both the crude (OR = 0.89, 95% CI: 0.84–0.93) and adjusted models (OR = 0.86, 95% CI: 0.82–0.91). Same trends were also observed in sensitivity analysis in both the crude (OR = 0.88, 95% CI: 0.81–0.97) and adjusted models (OR = 0.85, 95% CI: 0.77–0.94).

In the categorical variable evaluation, as compared with the participants with albumin levels of ≥4.7 g/dL, the risk of abnormal decline in kidney function was significantly higher not only in those with albumin levels of ≤4.3 g/dL but also in those with 4.4-4.6 g/dL (Fig. 3A): This result means that lower serum albumin level, even within the normal range, is related to rapid kidney function decline. The results of the sensitivity analysis, in which a ΔeGFR of ≥5 mL/min/1.73 m^2^/year was regarded as an abnormal decline in kidney function, are shown in Fig. 3B: Similar trends were observed.

**Figure 3.**
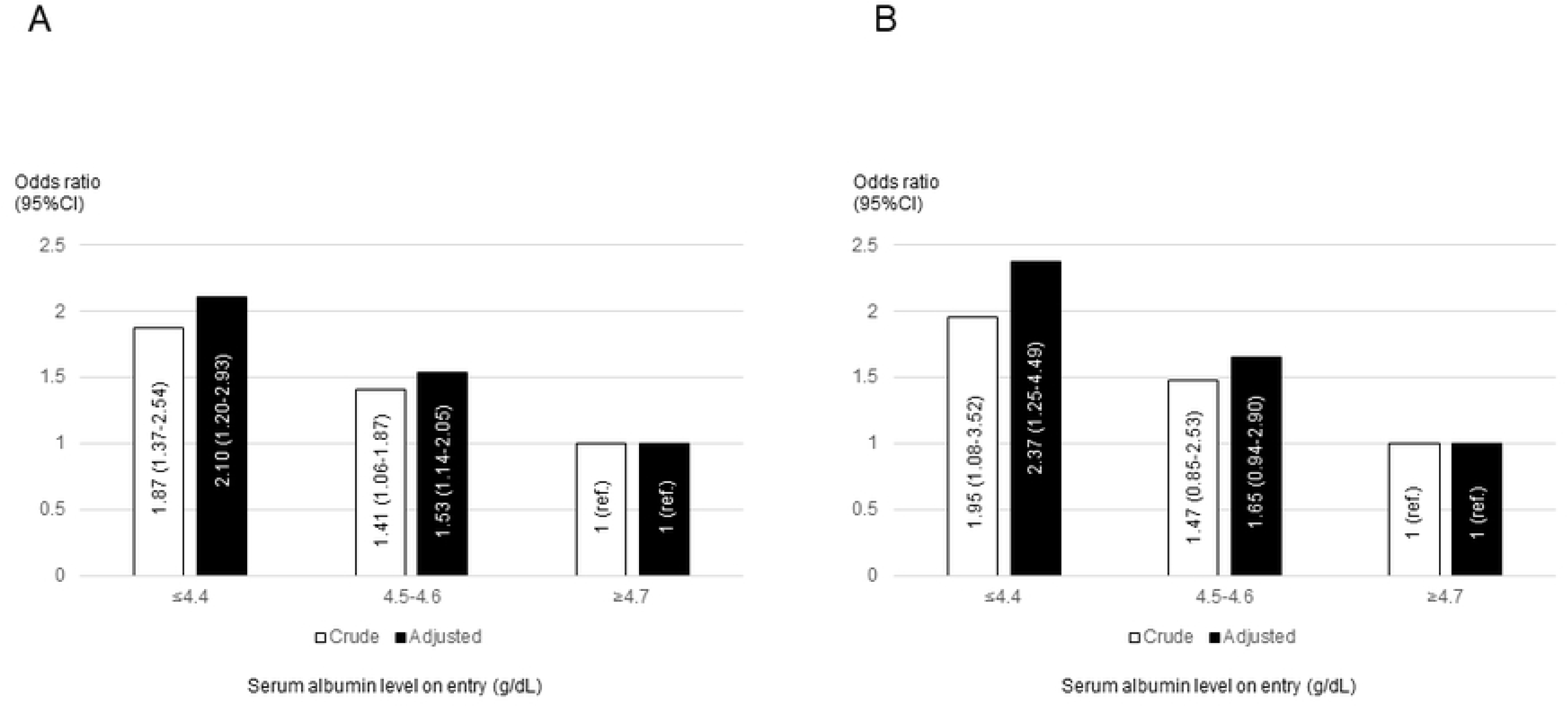
Association between albumin level at baseline and renal outcome. A: main analysis in which renal outcome is ΔeGFR/year ≥3 mL/min/1.73 m^2^, B: sensitivity analysis in which renal outcome is ΔeGFR/year ≥5 mL/min/1.73 m^2^. In both analyses, similar trends were observed: Namely, the risk of kidney impairment was increased in correlation with lowering of serum albumin even within normal range (P-value; <0.0001 in A and =0.0249 in B, Cochran-Armitage test). Adjusted for age, sex, body mass index, systolic blood pressure, eGFR at baseline, serum alanine aminotransferase level, serum uric acid, high-density lipoprotein cholesterol, HbA1c level, C-reactive protein level, history of cardiovascular disease. Abbreviations: CI, confidence interval; eGFR, estimated glomerular filtration.

## DISCUSSION

In this retrospective, population-based, cohort study, the relationship between albumin level and decline in kidney function over time was investigated. A decrease in albumin level was found to be the primary risk factor of abnormal decline in kidney function. More importantly, our study suggests that even within the normal range, those with albumin levels of ≤4.6 g/dL had a risk of decline in kidney function.

Our study could not reveal the reason why relatively low but within normal limit level of albumin read to early decline of kidney function. Slight decline of serum albumin might reflect some confounding risk factor such as slight undernutrition [24], slight albuminuria, or profound liver dysfunction. Another explanation is that relatively lower level of serum albumin *per se* caused rapid decline of kidney function. Serum albumin, which epithet is “multi-functional protein” [25], has various functions as follows: maintenance of osmotic pressure, buffering of the acid-base balance, supply of amino acid to tissues, binding and transporting of numerous compounds, elastase activity, and antioxidative activity. In fact, serum albumin level is the most abundant, thus most important, antioxidant of the extracellular space [26]. We reported that the decrease in serum albumin fraction, which has an antioxidative property, correlated with kidney dysfunction [27, 28] and that such decrease in serum albumin fraction was directly related to cardiovascular incidence in the population with advanced CKD [29, 30]. Therefore, theoretically and actually, lower serum albumin level can induce undesirable outcomes such as mortality [14], cardiovascular incidence [15], and rapid decline in kidney function as shown in the present study.

Most previous studies used abnormally lower albumin levels as an indicator of malnutrition status for research [15,31]. However, our study showed that even within the normal range, albumin levels of ≤4.6 g/dL are associated with a risk of decline in kidney function. Only a few studies showed that lower albumin levels within the normal range affect kidney function. Among 2,535 subjects aged 40–69 years, in the multivariable analyses, the CKD hazard ratios (95% CI) for the highest and lowest quartiles of serum albumin levels were 0.69 (0.40– 1.17) for men and 0.42 (0.28–0.64) for women [32]. While these results were similar to ours, the number of subjects was small. To overcome this limitation, in our study, we included almost five times more subjects than that in the previous study.

Our study showed that decreased albumin level was significantly associated with abnormal decline in kidney function and patients with albumin levels of ≤4.6 g/dL had a risk of decline in kidney function. This indicated that a slight decrease within the normal range of albumin level might be a risk for kidney function deterioration. This is consistent with the results of a previous study that reported that lower albumin level within the normal range was predictive of CKD in women [32]. Together with a previous observational study, our findings suggested that the risk of reduced kidney function might increase with decreased albumin levels even within the normal range. These findings might be therapeutic target from the viewpoint of public health.

This study has some limitations. First, the findings cannot be generalized to other ethnic or age groups, as the study participants might be healthier than the general population and their risk of developing medical complications was lower, as the study subjects were participants in a health checkup program. Second, the albumin levels were measured only at baseline; the changes in albumin levels during the follow up period that might have an independent effect on kidney outcome were not evaluated. Third, the eGFR was evaluated only twice; this parameter is known to show day-to-day variations. Fourth, present study lacks data regarding urinary finding: the level of urinary albumin might relate to the serum albumin level. Lastly, further studies are needed to assess whether other confounders such as smoking history and serum calcium/phosphate levels may affect kidney function. Future prospective studies on these relationships should be conducted to provide more insight.

In conclusion, our study showed that decreased serum albumin level is an independent risk factor of abnormal decline in kidney function in the general population and that a slight decrease in albumin level, even within the normal range, may be a risk factor of decline in kidney function.

